# S1000: A better taxonomic name corpus for biomedical information extraction

**DOI:** 10.1101/2023.02.20.528934

**Authors:** Jouni Luoma, Katerina Nastou, Tomoko Ohta, Harttu Toivonen, Evangelos Pafilis, Lars Juhl Jensen, Sampo Pyysalo

## Abstract

**Motivation:** The recognition of mentions of species names in text is a critically important task for biomedical text mining. While deep learning-based methods have made great advances in many named entity recognition tasks, results for species name recognition remain poor. We hypothesize that this is primarily due to the lack of appropriate corpora.

**Results:** We introduce the S1000 corpus, a comprehensive manual re-annotation and extension of the S800 corpus. We demonstrate that S1000 makes highly accurate recognition of species names possible (F-score=93.1%), both for deep learning and dictionary-based methods.

**Availability:** All resources introduced in this study are available under open licenses from https://jensenlab.org/resources/s1000/. The webpage contains links to a Zenodo project and two GitHub repositories associated with the study.

**Supplementary information:** Supplementary data are available at *Bioinformatics* online.

## 1 Introduction

Over the last two decades of research into information extraction from biomedical scientific publications, progress has primarily been driven by advances in two key areas: manually annotated corpora, and deep learning methodology. Corpora of sufficient size, coverage and annotation quality have been established to allow the development of methods capable of highly accurate recognition of many key entity types, including simple chemicals (Krallinger *et al.*, 2015; Li *et al.*, 2016), genes and proteins (Kim *et al.*, 2004; Smith *et al.*, 2008), diseases (Doğan *et al.*, 2014) and anatomical entities (Pyysalo and Ananiadou, 2014). Using these resources, state-of-the-art methods approach or exceed 90% precision or recall at the recognition of mentions of the names of these entities (Lee *et al.*, 2019; Lewis *et al.*, 2020; Shin *et al.*, 2020). However, strikingly, these same methods fail to achieve comparable levels of performance in the recognition of species names, a highly relevant target for biomedical information extraction that one would intuitively expect to be comparatively simple to recognize due to the regularity of the binomial nomenclature and the availability of high-coverage resources of species names (Schoch *et al.*, 2020).

Most recent efforts targeting species name recognition focus on two manually annotated resources: the LINNAEUS corpus (Gerner *et al.*, 2010) and the Species-800 (S800) corpus (Pafilis *et al.*, 2013). The LINNEAUS corpus of 100 full-text articles was the first big effort to generate a manually annotated corpus for evaluating Named Entity Recognition (NER) and normalization for species. Following LINNEAUS, the S800 corpus aimed at increasing the diversity of species annotations and coverage of different life domains in comparison to the former. For these reasons a corpus was compiled consisting of abstracts – instead of full-text documents – from 2011 and 2012, published in journals representing to eight different categories: seven positive categories for different taxonomic groups and an eighth category (Medicine) primarily included as a negative control.

Both of these corpora were originally introduced to support the development and evaluation of dictionary-based NER tools. Existing tools use dictionaries based on NCBI Taxonomy (Schoch *et al.*, 2020), which includes e.g. subspecies and strains in addition to species. These tools generally focused on identifying the right entities in text rather than on getting the species name boundaries perfectly right. Evaluations were thus done using relaxed boundary matching and corpus annotation guidelines consequently did not need to have stringent definitions for how to annotate boundaries. While this is no problem for the originally intended purpose of these corpora, most recent Deep Learning (DL) experiments on them (Zhang *et al.*, 2021; Phan *et al.*, 2021; Kocaman and Talby, 2021; Lewis *et al.*, 2020; Lee *et al.*, 2019; Sharma and Daniel Jr, 2019; Giorgi and Bader, 2018) use the standard exact matching criteria established in the CoNLL shared tasks, which then becomes problematic.

In fact, currently no corpus exists that is well-designed for DL-based species detection. Most corpora, including LINNAEUS and broader corpora like GENIA (Kim *et al.*, 2003), have been created in ways that resulted in human and model organisms making up the vast majority of the species annotations. This low diversity of species in most corpora can already be an issue during evaluation of DL-based methods, but is much more problematic when the corpora are used for training. For the purpose of training DL-based methods to recognize species names, ideally a corpus would contain a wide range of different species names covering all kingdoms of life, systematically annotated with accurate boundaries regardless of whether the species is included in NCBI Taxonomy.

Here we present S1000, a corpus for species NER, which builds upon S800. S800 was chosen as a starting point, since it already fulfills the criteria of species name diversity and representation. Several improvements were made so that S1000 can support training of state-of-the-art DL-based methods for species NER, while at the same time continue serving its original purpose of a corpus for the evaluation of dictionary-based NER methods. We present herein that the use of S1000 leads to a significant increase in performance, when used in training DL-based methods, while at the same time the addition of 200 documents increases its generalizability. We are certain that S1000 can replace S800, and will be the new gold standard for evaluating DL-based methods.

## 2 Materials and Methods

### 2.1 Manual revision of corpus annotation

The revision of the corpus annotation consisted of the following primary steps:

- Decoupling of recognition from normalization (***Decoupling***)
- Revision of annotations for boundary consistency (***Boundary consistency***)
- Separation of strain from species mentions (***Strains***)
- Annotation of genera (***Genera***)
- Extension of corpus with additional documents and final polish (***Extension***)

In the following, we briefly describe these steps. More details regarding the annotation rules followed to produce the corpus can be found in the annotation documentation that the annotators have used ^1^.

#### 2.1.1 Decoupling of recognition from normalization

The original S800 corpus only annotated species mentions that could be normalized to a version of the NCBI Taxonomy from 2013 (Pafilis *et al.*, 2013). This made sense from the standpoint of evaluating dictionary-based methods developed at the time, since none of them would be able to recognize and normalize species not existing in NCBI Taxonomy. However, from the perspective of pure NER of species names (regardless of normalization), this caused the annotation to appear incomplete in places. In the first revision step, annotation was added for scientific and common names of species regardless of whether they could be normalized to an NCBI Taxonomy identifier in the version of the database published in 2020 (Schoch *et al.*, 2020). A revision pass addressing the overall consistency of annotation was performed, and annotated names of genera, families and other levels of taxonomy above species were annotated as “out-of-scope” during this process. Moreover, genus or higher-level mentions (e.g. Arabidopsis, yeast) that were originally annotated as synonyms of species names, received an annotation corresponding to their real taxonomic level (e.g. genus for Arabidopsis). Annotated entities include only taxonomic and common names, which means that nominal non-name “species clues”, such as “patient” or “woman” — which are annotated in the LINNEAUS corpus (Gerner *et al.*, 2010), but not in the original S800 corpus — remained unannotated.

#### 2.1.2 Revision of annotations for boundary consistency

The original evaluation of taggers using the S800 corpus (Pafilis *et al.*, 2013) applied relaxed boundary matching criteria. As a result, any tag that overlapped with a sub-string of a manually annotated species entity with the correct taxonomic identifier assigned was regarded as a true positive for evaluation purposes. This resulted in the boundaries of annotated mentions to be inconsistently annotated in many places, which as explained above, is a problem when training machine learning-based methods. To address this issue, we created detailed guidelines on how to determine entity boundaries and made a revision pass addressing span consistency issues in the data set. This revision step also included a focused review of the annotation of virus mentions, which had comparatively frequent annotation boundary issues. During this revision step, organism mentions of taxonomic rank genus and above in the “Viruses” superkingdom were corrected to better reflect their place in the lineage, thus fixing cases of imprecise normalization to species mentions in the original S800 corpus.

#### 2.1.3 Separation of strain mentions from species mentions

The original S800 corpus annotation only involves a single annotated mention type (“species”) that is used to annotate mentions of species names, as well as mentions of strains. In this revision of the corpus, we introduced a separate “strain” type and revised all strain name mentions to use this type, also revising the spans of species annotations to exclude strain names when the two occurred together in text. The lack of a universally accepted definition of “strain”, both within specific communities (e.g. virology (Kuhn *et al.*, 2013)) and across different communities, makes the strict definition of “strain” entity type impossible. For this reason we decided, to also annotate other fine-grained taxa from NCBI Taxonomy (Schoch *et al.*, 2020), such as isolates and serotypes, as “strain” in the corpus. The only taxonomic groups “below species level” that were treated differently are “subspecies” and “biotypes”, where entire mentions were simply annotated as slightly longer “species” mentions. Finally, cultivars and ecotypes, which are not a not official taxonomic ranks, were treated as “out-of-scope”.

#### 2.1.4 Annotation of genera

The original S800 corpus annotation included partial annotation for mentions of names at taxonomic ranks above species, in particular in a number of cases where these names were used in an imprecise way to refer to species (e.g. Drosophila for *Drosophila melanogaster).* As mentioned above, during the initial revision step addressing annotation consistency, we marked such cases as “out-of-scope”. This resulted in the reduction of the coverage of the revised annotation in some aspects from that of the original S800 corpus. To partially remedy this issue, we reintroduced annotation for mentions of names at the “genus” taxonomic rank in a systematic way, by adding it as a distinct annotated type. For annotations above the “species” rank only the “coarse” ranks have been considered, thus mapping mentions at fine-grained levels to their coarse equivalents, e.g. “subgenus” maps to “genus”. “Clade” rank is a new monophyletic, non-hierarchical rank, introduced in the latest revision of NCBI taxonomy. Since this rank is non-hierarchical, and can appear anywhere in the lineage without breaking the order, mentions normalizing to NCBI taxonomy nodes with the “clade” rank have been assigned the type based on the rank of their first non-clade ancestor node, when that was in scope of our annotation.

#### 2.1.5 Extension of corpus with additional documents and final polish

Despite its efforts to increase diversity, there is still a concern that since all the articles in S800 originate from the same journals, any method trained using this data might overfit to that specific corpus during development and might not perform as well in an open-domain annotation task. To alleviate this concern we decided to introduce even more diversity in the corpus, via its extension with 200 additional documents, thus generating the extended version of S800, called S1000. The selection process was such so that the new documents would not be limited to specific preselected journals and specific publication years. In the next two subsections the selection process of documents in the positive and negative categories is explained in more detail.

##### Positive categories

For the extension of abstracts in the positive categories, we wanted to select publications that would contain species mentions, whose genera are not represented in the original S800 corpus. This includes all the genera of the species in S800, retrieved by mapping the species mentions to their parental rank in NCBI Taxonomy. To avoid biasing the new documents towards what can be found by a specific text-mining system, we decided not to use text mining to find candidate abstracts. Instead, we used the literature references within manually curated UniProtKB/SwissProt (SwissProt hereafter), which are added by annotators as the primary source to support annotations of proteins (The Uniprot Consortium, 2021). This allows easy automatic retrieval of the corresponding abstracts, which will commonly mention the name of the species that the protein is from, despite the fact that SwissProt is a protein resource. Thus, we could use this strategy to obtain a broad selection of candidate abstracts containing species names from genera complementary to those already included in S800.

An advantage of the original S800 over other corpora, is the fact that documents come from different categories, which in turn allows better performance evaluation during benchmarking, as it can be assessed whether a method is better “across the board” or in a specific domain. This is a property we wanted to maintain during the corpus extension in S1000. An added advantage of using SwissProt to detect candidate documents for annotation, is that it permits the placement of documents in the same categories as in the original corpus, which both retains balance and allows the new process to be consistent with what was originally done. Specifically, since SwissProt entries have information about the taxonomy of the species to which a protein belongs to, we decided to map the document categories of the original S800, to taxonomic ranks in NCBI taxonomy. This allows to label documents as belonging to a specific category based on the taxonomy of the species a protein belongs to. The mapping and the number of documents for each category are shown in Table 1. Finally, 25 documents were randomly selected for each of the positive categories and were added to the corpus.

**Table 1.**
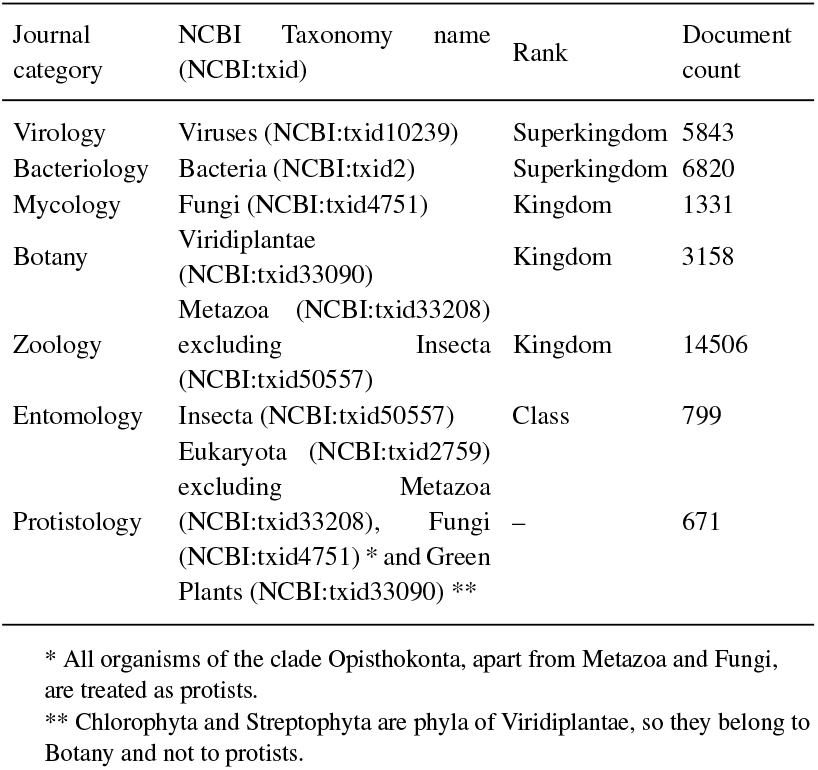
Mapping between categories in S800 and NCBI Taxonomy. The number of documents found in SwissProt for proteins belonging to organisms in each of these taxonomic groups is provided.

##### Negative category

In addition to the categories mentioned in Table 1, the original S800 contained 100 abstracts from the medical literature, which served as a negative category in which not many species’ names mentions were expected to show up. To detect documents that would serve as a negative control for S1000, but to avoid focusing on specific journals or publication years, we aimed our selection towards PubMed abstracts where a species tagger (Pafilis *et al.,* 2013) had not detected any species mentions. In total, there were 20,320,693 documents with no species mentions detected, and from those we randomly selected 25 to form the negative category for the S1000 corpus.

As a final step, a semi-automated check was performed to evaluate the consistency of mentions in text and the names and synonyms of the NCBI taxonomy entries that these normalize to. This produced a list of common and scientific species and genus names that do not have a clear match in the NCBI taxonomy. All these names were manually checked against alternative taxonomic resources (namely ITIS^2^, Catalogue of Life^3^, Avibase^4^, ICTV^5^ and WoRMS^6^) to assess whether a link to NCBI taxonomy entries could be obtained via them. Where possible, species and genera synonyms were added for mapping between the surface form and the taxonomy name.

### 2.2 Dictionary-based NER

The JensenLab tagger^7^ (Jensen, 2016) is a dictionary-based method used for the recognition of species mentions, among other biomedical entities. The species NER is extremely important for the text-mining evidence channel in the influential database of protein–protein interactions STRING (Szklarczyk *et al.*, 2023), as it allows both the recognition of the species of origin for proteins mentioned in text, as well as the disambiguation of ambiguous protein names, based on species mentions in the document. It is also important for other resources, like ORGANISMS (Pafilis *et al.*, 2013) with tagging results of organism names in the scientific literature.

As already mentioned, biomedical corpora for species names, like S800, had been originally developed with the purpose of evaluating dictionary-based methods. To make sure that the revised version of the corpus is still suitable for this original purpose, the final revised annotation of S1000 was used to evaluate JensenLab tagger. This evaluation focused solely on mentions of type “species” in the corpus. The JensenLab tagger software was run on the S1000 corpus test set and taxonomic identifiers were mapped to their corresponding taxa. All identifiers above species level were ignored and all mentions of taxa below species level were assigned to their parent species and were kept during this evaluation. This was done for consistency with how the JensenLab tagger works, as it uses the taxonomy structure to backtrack names at lower taxonomic levels to all their parent levels. Moreover, mentions in two branches of the NCBI taxonomy – namely “other entries” and “unclassified entries” – which contain metagenomes, plasmids and other similar entries, were out of scope for the annotation effort of S1000 and were also ignored during the evaluation phase.

### 2.3 Transformer-based

Since the majority of the biomedical text-mining community has now migrated to DL-based and specifically Transformer-based methods — as shown e.g. in Miranda *et al.* (2021) — we needed to make sure that this corpus can serve the purposes of both training and evaluating DL-based methods. The current state-of-the-art methods in NER dominantly utilize models based on the Transformer architecture (Vaswani *et al.*, 2017), and for that reason we focused our efforts on these. These models are initially pre-trained on large collections of text to produce a general language model. Such models can then be fine-tuned to perform specific tasks such as NER. We have selected three pre-trained models for closer evaluation, namely RoBERTa-large-PM-M3-Voc (hereafter RoBERTa-biolm) (Lewis *et al.,* 2020), BioBERTLarge, cased (hereafterBioBERT) (Lee *et al.,* 2019) and BioMegatron 345M Bio-vocab-50k, case (hereafter BioMegatron) (Shin *et al.*, 2020). These models have been pre-trained on biomedical literature and have shown good performance in NER tasks for biomedical texts.

### 2.4 Experimental setting

We used the method proposed in (Luoma and Pyysalo, 2020) for training and evaluation of the Transformer-based models. We fine-tuned the models to detect the available entity types in the training data (“species”, “strain” and “genus” depending on the corpus’ revision step) with attaching a single fully connected layer on top of the Transformer architecture for classifying individual tokens in input samples.

The training and evaluation of all of the steps except the last (***Expansion***) are done on the same original documents that created the S800 corpus. Initially the documents were split to separate training, development and test sets: 560 documents for training, 80 for development, and 160 for test set, using the standard split introduced in Hakala *et al.* (2016). The final stage of extending the corpus brings the numbers up to 700 documents for training, 100 for development and 200 for the test set, while still respecting the original split. For this research the training and development sets were combined and then split to eight folds with stratification over the original publication sources. The folds on document level were kept the same for each corpus development step with documents added on each fold in the last step (***Expansion***).

The hyperparameter selection was done using a grid search. The experiments with each combination of hyperparameters were run in a cross-validation setup to reduce the effects of over-fitting to development set and to reduce the effect of random events in the training process (e.g. layer initialization, dropout) on the results. The hyperparameters producing the best total mean F1-score on the 8-fold cross-validation for all of the mention types was selected for training the models for evaluation on test set. For the final evaluation against the test set, all training and development data was used in fine-tuning the models with the optimal hyperparameters. The process was repeated five times and the results are expressed as a mean and standard deviation of the exact match F1-score (micro-averaged over all mention types).

We first compared the performance of the three different Transformer-based models on the whole S1000 corpus and then selected the best performing model as basis for further evaluation of the performance on different corpus annotation steps.

The progression of the performance on different corpus revision steps defined in the section 2.1 was concentrating on species mentions. In addition to the evaluation on exact match F1-score, we evaluated the single model reaching the best exact match F1 score on test data using the overlapping matching criterion, to compare the performance with the dictionary-based tagger.

## 3 Results and Discussion

### 3.1 Corpus statistics

In Table 2 an overview of corpus statistics is shown. The numbers for unique names and total mentions of entities belonging to all taxonomic levels that were in scope for this annotation effort are shown. In this reannotation and extension effort we have almost doubled the number of unique and total mentions compared to the S800 corpus. This is of course mostly due to extending the scope from only “species” in S800 to include also “strains” and “genera” in S1000. But even the number of “species” alone has increased by 21% (from 3708 to 4506), and the number of unique “species” names by 16% (from 1503 to 1756).

**Table 2.** S1000 corpus statistics. The numbers for unique and total mentions for S800 as presented in the original publication (Pafilis *et al.*, 2013) are also provided.

| Category | Unique Names |  | Total Mentions |  |
| --- | --- | --- | --- | --- |
|  | S1000 | S800 | S1000 | S800 |
| Bacteriology | 316 | 179 | 788 | 416 |
| Botany | 217 | 131 | 510 | 308 |
| Entomology | 441 | 293 | 965 | 614 |
| Mycology | 266 | 178 | 784 | 538 |
| Protistology | 638 | 284 | 1104 | 497 |
| Virology | 532 | 342 | 1539 | 946 |
| Zoology | 283 | 160 | 519 | 299 |
| Medicine | 46 | 30 | 119 | 90 |
| Total | 2583 | 1503 | 6328 | 3708 |

In a broader context, the S1000 corpus contains more than seven times as many unique names as the LINNAEUS corpus (2583/375). The high diversity of names was one of the key motivators for choosing S800 as a starting point, and our efforts to increase it even more have paid off, as is clear from the corpus statistics presented in Table 2.

More detailed corpus statistics are available in *Supplementary Table* 1.

### 3.2 Evaluation on Transformer-based models

We used the combined training and development sets of the S1000 corpus to fine-tune different pre-trained Transformer-based models and evaluated their performance against the test set of the corpus. The results of these tests are expressed as mean entity level exact match F1-scores and standard deviations of five repetitions of the test. The results are shown in Table 3. The numbers are consistently around 90% for total F1-score and over 90% for “species” mentions for all of the tested models, showcasing even further that the S1000 corpus provides improvements in recognition of “species” mentions when compared to the earlier S800 corpus.

**Table 3.** Model comparison on S1000. Results are presented for all mention types on the test set. The exact matching criterion is used during this evaluation.

| Type | RoBERTa-biolm |  | BioMegatron |  | BioBERT |  |
| --- | --- | --- | --- | --- | --- | --- |
|  | F1 | std | F1 | std | F1 | std |
| Species | **93.14** | 0.79 | 91.48 | 0.57 | 91.20 | 0.42 |
| Genus | **91.45** | 0.72 | 86.52 | 1.72 | 88.44 | 1.40 |
| Strain | 80.28 | 1.58 | **80.61** | 1.06 | 78.75 | 2.94 |
| Total | **91.07** | 0.69 | 89.23 | 0.47 | 89.04 | 0.52 |

\*F1: F1-score, std: standard deviation

We find that the RoBERTa-biolm model outperforms the other two on this data set, partially agreeing with the findings of Lewis *et al.* (2020), where the RoBERTa-biolm was found performing better than BioBERT on various biomedical NER datasets.

For detailed results please refer to *Supplementary Table 2.*

### 3.3 Progression

The progression of the results in tagging performance of “species” mentions with RoBERTa-biolm are shown in Figure 1. The **exact matching** criterion is used during this evaluation. From the figure it can be seen that each of the first three corpus revision steps increases the performance on “species” mentions. Then the addition of “genus” mentions causes a slight decrease in recall, but continues to improve precision. Finally the addition of 200 documents with more diverse names causes a decrease in performance, reflecting the new challenges introduced by the addition of documents, as originally intended. Specifically, the extension can actually help generate a corpus with higher variance, which in turn allows the training of more “generalizable” models. At the same time it does not take away from the huge progress that was made from the initial to the final revision step, making – to the best of our knowledge – S1000 the best DL-ready corpus for species NER currently in existence.

**Fig. 1.**
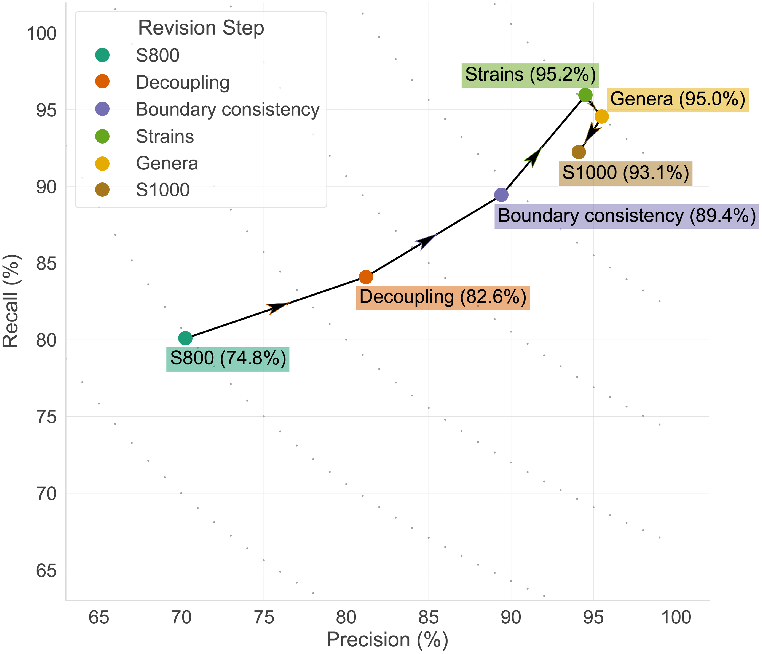
Performance on species mentions on test data for different corpus revision steps. The arrows denote the sequence of the progression. In parentheses next to each step the F-score is provided. The different corpus revision steps are defined in Section 2.1.

For detailed results please refer to *Supplementary Table 3.*

### 3.4 Performance comparison and error analysis

#### 3.4.1 Evaluation on the entire corpus

From the progression curve presented above (Section 3.3) it is clear that S1000 is a better corpus for training DL-based methods. But we also needed to test if it can serve its original purpose of evaluating dictionary-based methods. To test this, we applied both the dictionary-based (Jensenlab) and Transformer-based taggers on the S1000 test set and evaluated them on “species” names detection. Since the JensenLab tagger finds left-most longest matches of the names in its dictionary, which includes more than just species names, the **overlapping matching** criterion is used to evaluate both methods. The F-score for dictionary-based tagger on this set is 84.7% (precision: 87.3%, recall: 82.3%), while for the Transformer-based tagger is 97.0% (precision: 97.9%, recall: 96.1%). The results for the Transformer-based model are even better than those reported above, since the switch from using the exact to the overlapping matching criterion during evaluation, eliminates all boundary inconsistency errors for this method.

An analysis of the errors produced by both methods, grouped in categories, is presented in Table 4. For a detailed overview of all the errors, please refer to *Supplementary Tables 4* and 5, for the JensenLab tagger and the Transformer-based tagger, respectively.

**Table 4.** Error analysis for the JensenLab tagger and the Transformer-based tagger.

| Error categories | Total |  | FN |  | FP |  |
| --- | --- | --- | --- | --- | --- | --- |
|  | dict-based | TF-based | dict-based | TF-based | dict-based | TF-based |
| Ambiguous name | 35 | 18 | 34 | 10 | 1 | 8 |
| Annotation error | 9 | 2 | 7 | 0 | 2 | 2 |
| Dictionary error | 157 | – | 113 | – | 44 | – |
| Discontinuous entity | 6 | 0 | 6 | 0 | 0 | 0 |
| Lower taxonomic level tagged as species | 56 | 0 | 0 | 0 | 56 | 0 |
| Upper taxonomic level tagged as species | 17 | 10 | 0 | 0 | 17 | 10 |
| Transformer-based model error | – | 24 | – | 24 | – | 0 |
| Unofficial abbreviation | 12 | 4 | 12 | 4 | 0 | 0 |
| Total | 292 | 58 | 172 | 38 | 120 | 20 |

\*FN: false negative, FP: false positive, dict-based: dictionary-based tagger, TF-based:Transformer-based tagger

More than 50% of the errors produced by the dictionary-based tagger are “dictionary errors”. As is evident by their name, dictionary-based methods are only as good as their source dictionary, and this inherent property is clearly reflected in the error analysis performed above. Most of the dictionary errors affect its recall, meaning that the issue we observe here is mostly names missing from the dictionary. “Discontinuous entities” and “unofficial abbreviations” are another issue mainly faced by the dictionary-based method. Even though, these are clearly problems that affect the dictionary-based method in this analysis, it should be noted that all these problems would also affect the Transformer-based method if normalization was done on top of recognition, and NCBI taxonomy was used as the source for this normalization. There are of course measures one could take to reduce such issues (e.g. using similarity metrics to identify synonyms and resolve unofficial abbreviations), but when names are completely missing from the source used for normalization, there is little one can do.

Both methods seem to have issues with “ambiguous names”, like *bees*, which can be both a species and a superfamily name. Considering the nature of such ambiguity – i.e. the fact that the ambiguous name is a taxonomic name in either case – it is easy to understand why such entities could affect the performance of either method. “Upper taxonomic level tagged as species” was a problem that affected both methods similarly. For the dictionary-based tagger these errors could be actually counted as dictionary errors, since they reflect errors in synonyms assigned to “species” instead of upper taxonomic ranks in NCBI taxonomy. For the Transformer-based model errors of this type and “Transformer-based model error” are a result of “clade” or “no rank” entities in NCBI taxonomy being annotated as “species” (see Section 2.1.4 for more details) and common organism names either not being detected, or being misclassified in regards to their position in the taxonomy.

As mentioned in the Methods section, the JensenLab tagger always backtracks lower taxonomic level mentions to their “species” parent. This leads to one type of error in Table 4, “lower taxonomic level tagged as species”, which affects only the dictionary-based method and might as well not be considered actual errors. If these errors did not count as False Negatives then the True Positive count would increase from 811 to 867. Similarly, if “annotation errors” were not counted as either False Negatives or False Positives, then the True Negative count would decrease from 984 to 982 and the True Positive count would increase to 874. If metrics are then recalculated for the dictionary-based method, the precision would now be 94.1%, the recall 82.5% and the F-score 87.9%. Not counting “annotation errors” for the Transformer-based tagger would also slightly increase its precision to 98.13% and F-score to 97.12%. The dictionary-based tagger seems to perform much better, if these errors are not counted, but still cannot outperform the Transformer-based model, since the majority of the errors continues to be due to shortcomings of the dictionary, as already discussed above.

#### 3.4.2 Evaluation per journal category

The design of the S800 corpus, and consequently also of S1000, allows us to delve deeper when assessing the errors produced by both dictionary and DL-based methods. S1000 consists of eight journal categories, corresponding to seven taxonomic groups (see Table 1) and a negative class. These can be used to identify whether specific categories of documents – and as a consequence specific parts of the taxonomy – are more difficult to detect in text.

The Transformer-based method (Fig.2, triangles) seems to perform consistently well in all journal categories, with both precision and recall over 95% for all categories, except Virology. The recall for the dictionary-based tagger is consistently lower, but as explained in Section 3.4.1 this is mostly due to names missing from the dictionary. To better assess what explains the differences in recall, one can examine a journal category where the precision for the two methods is similar, but the recall is significantly different, like Zoology (Fig. 2, yellow). When one examines the errors in *Supplementary Tables 4* and *5*, it is obvious that both methods have a problem with “gibbon” which is tagged as a “species” mention instead of “family” for both. All the remaining errors, that lower the recall for the dictionary-based method, are cases of common species names that are missing as synonyms for the respective NCBI taxonomy entries. When examining another category with differences both in precision and recall, like Bacteriology (Fig. 2, blue), it seems that the vast majority of false positives are names that should be blocked, while the false negatives are Linnean names that are missing from NCBI taxonomy. In this category it is obvious that the precision of the dictionary-based method could be improved, e.g. by using a deep-learning method to suggest names to block from tagging. This is a project that we are currently working on to improve our dictionary. As previously noted, False Negatives mainly occur due to the absence of certain common and Linnean taxonomic names in NCBI Taxonomy.

**Fig. 2.**
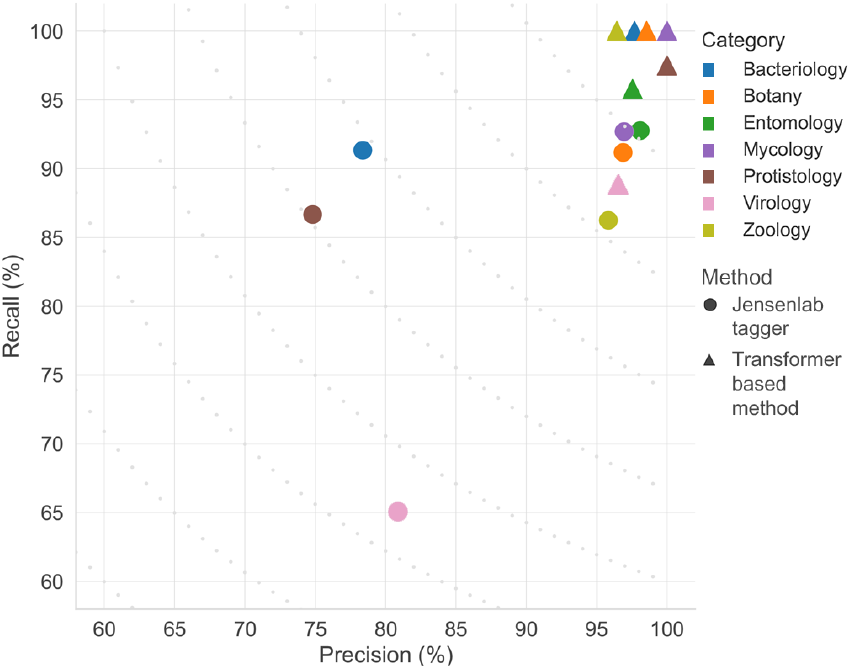
Precision-Recall plot for the dictionary-based and the Transformer-based taggers on the seven S1000 journal categories. F-score contours are presented with grey dots in the plot.

Both methods perform worst on papers from Virology journals. This is probably due to the non-systematic naming conventions for viruses and the extensive use of acronyms in this field, which result in names that are difficult to capture for both DL- and dictionary-based methods. The error analysis for the dictionary-based method shows that the two main sources of errors in this journal category are once again either entities missing from the dictionary or lower taxonomic level entities captured as species (with the latter not being a problem in a real-world scenario, as explained in Section 3.4.1).

### 3.5 Large-scale tagging

Results on tagging of PubMed abstracts (as of August 2022) and articles from the PMC open access subset (as of April 2022) for both the JensenLab tagger and the Transformer-based method are provided via Zenodo^8^. There are in total 185,869,193 organism matches for JensenLab tagger, amongst which the vast majority (176,700,642) are species or subspecies mentions backtracked to species, covering 818,547 unique species names. Tagging with the Transformer-based model yielded 196,511,523 total matches, comprising 142,522,111 species, 36,461,427 strain and 17,527,985 genus matches. These are comprised of 4,041,604 unique names, of which 1,953,694 are species.

## 4 Conclusions

In this work we present S1000, a re-annotated and expanded high-quality corpus for species, strain and genera names. We propose the use of this improved corpus as a gold standard for the evaluation of deep learning-based language models in the place of the already widely used S800 corpus. Our experiments have shown that the use of S1000 results in a clear improvement in performance with an 18.3% increase in F-score (from 74.8% in S800 to 93.1% in S1000). This was achieved mainly because the re-annotation effort focused on ensuring that S1000 can support the training of state-of-the-art DL-based models. Moreover, the expansion with 200 additional documents allows training of more generalizable models, while still maintaining over 90% F-score. We have also demonstrated that the annotation improvements have not affected our ability to use the corpus for the evaluation of dictionary-based NER methods, on top of DL-based methods. Notably, the unique and total mentions of names in S1000 has almost doubled in comparison to S800. Finally, all data used in this project, along with the code to reproduce the results, are publicly available, including results of large-scale tagging of the entire literature.

## Acknowledgements

We thank the CSC – IT Center for Science for generous computational resources.

## Funding

This project has received funding from Novo Nordisk Foundation (Grant no.: NNF14CC0001) and from the Academy of Finland (Grant no.: 332844). K.N. has received funding from the European Union’s Horizon 2020 research and innovation programme under the Marie Sklodowska-Curie (Grant no.: 101023676).

## Footnotes

1 https://katnastou.github.io/s1000-corpus-annotation-guidelines

2 https://www.itis.gov/

3 https://www.catalogueoflife.org/

4 https://avibase.bsc-eoc.org/

5 https://talk.ictvonline.org/taxonomy/

6 https://www.marinespecies.org/

7 https://github.com/larsjuhljensen/tagger

8 https://zenodo.org/deposit/7064902

## Notes

https://jensenlab.org/resources/s1000/

### Competing Interest Statement

The authors have declared no competing interest.

## References

Doğan, R. I., Leaman, R., and Lu, Z. (2014). NCBI disease corpus: a resource for disease name recognition and concept normalization. Journal of biomedical informatics, 47, 1–10.

Gerner, M., Nenadic, G., and Bergman, C. M. (2010). LINNAEUS: a species name identification system for biomedical literature. BMC bioinformatics, 11(1), 85.

Giorgi, J. M. and Bader, G. D. (2018). Transfer learning for biomedical named entity recognition with neural networks. Bioinformatics, 34(23), 4087–4094.

Hakala, K., Kaewphan, S., Salakoski, T., and Ginter, F. (2016). Syntactic analyses and named entity recognition for pubmed and pubmed central—up-to-the-minute. In Proceedings of the 15th Workshop on Biomedical Natural Language Processing, pages 102–107.

Jensen, L. J. (2016). One tagger, many uses: Illustrating the power of ontologies in dictionary-based named entity recognition. bioRxiv, page 067132.

Kim, J.-D., Ohta, T., Tateisi, Y., and Tsujii, J. (2003). GENIA corpus—a semantically annotated corpus for bio-textmining. Bioinformatics, 19(suppl_1), i180–i182.

Kim, J.-D., Ohta, T., Tsuruoka, Y., Tateisi, Y., and Collier, N. (2004). Introduction to the bio-entity recognition task at jnlpba. In Proceedings of the international joint workshop on natural language processing in biomedicine and its applications, pages 70–75. Citeseer.

Kocaman, V. and Talby, D. (2021). Biomedical named entity recognition at scale. In A. Del Bimbo, R. Cucchiara, S. Sclaroff, G. M. Farinella, T. Mei, M. Bertini, H. J. Escalante, and R. Vezzani, editors, Pattern Recognition. ICPR International Workshops and Challenges, pages 635–646, Cham. Springer International Publishing.

Krallinger, M., Rabal, O., Leitner, F., Vazquez, M., Salgado, D., Lu, Z., Leaman, R., Lu, Y., Ji, D., Lowe, D. M., et al. (2015). The chemdner corpus of chemicals and drugs and its annotation principles. Journal of cheminformatics, 7(1), 1–17.

Kuhn, J. H., Bao, Y., Bavari, S., Becker, S., Bradfute, S., Brister, J. R., Bukreyev, A. A., Chandran, K., Davey, R. A., Dolnik, O., et al. (2013). Virus nomenclature below the species level: a standardized nomenclature for natural variants of viruses assigned to the family filoviridae. Archives of virology, 158(1), 301–311.

Lee, J., Yoon, W., Kim, S., Kim, D., Kim, S., So, C. H., and Kang, J. (2019). BioBERT: a pre-trained biomedical language representation model for biomedical text mining. Bioinformatics, 36(4), 1234–1240.

Lewis, P., Ott, M., Du, J., and Stoyanov, V. (2020). Pretrained language models for biomedical and clinical tasks: Understanding and extending the state-of-the-art. In Proceedings of the 3rd Clinical Natural Language Processing Workshop, pages 146–157, Online. Association for Computational Linguistics.

Li, J., Sun, Y., Johnson, R. J., Sciaky, D., Wei, C.-H., Leaman, R., Davis, A. P., Mattingly, C. J., Wiegers, T. C., and Lu, Z. (2016). Biocreative v cdr task corpus: a resource for chemical disease relation extraction. Database, 2016.

Luoma, J. and Pyysalo, S. (2020). Exploring cross-sentence contexts for named entity recognition with BERT. In Proceedings of the 28th International Conference on Computational Linguistics, pages 904–914, Barcelona, Spain (Online). International Committee on Computational Linguistics.

Miranda, A., Mehryary, F., Luoma, J., Pyysalo, S., Valencia, A., and Krallinger, M. (2021). Overview of drugprot biocreative vii track: quality evaluation and large scale text mining of drug-gene/protein relations. In Proceedings of the seventh BioCreative challenge evaluation workshop.

Pafilis, E., Frankild, S. P., Fanini, L., Faulwetter, S., Pavloudi, C., Vasileiadou, A., Arvanitidis, C., and Jensen, L. J. (2013). The species and organisms resources for fast and accurate identification of taxonomic names in text. PloS one, 8(6), e65390.

Phan, L. N., Anibal, J. T., Tran, H., Chanana, S., Bahadroglu, E., Peltekian, A., and Altan-Bonnet, G. (2021). Scifive: a text-to-text transformer model for biomedical literature.

Pyysalo, S. and Ananiadou, S. (2014). Anatomical entity mention recognition at literature scale. Bioinformatics, 30(6), 868–875.

Schoch, C. L., Ciufo, S., Domrachev, M., Hotton, C. L., Kannan, S., Khovanskaya, R., Leipe, D., Mcveigh, R., O’Neill, K., Robbertse, B., Sharma, S., Soussov, V., Sullivan, J. P., Sun, L., Turner, S., and Karsch-Mizrachi, I. (2020). NCBI Taxonomy: a comprehensive update on curation, resources and tools. Database, 2020. baaa062.

Sharma, S. and Daniel Jr, R. (2019). Bioflair: Pretrained pooled contextualized embeddings for biomedical sequence labeling tasks. arXiv preprint arXiv:1908.05760.

Shin, H.-C., Zhang, Y., Bakhturina, E., Puri, R., Patwary, M., Shoeybi, M., and Mani, R. (2020). BioMegatron: Larger biomedical domain language model. In Proceedings of the 2020 Conference on Empirical Methods in Natural Language Processing (EMNLP), pages 4700–4706, Online. Association for Computational Linguistics.

Smith, L., Tanabe, L. K., Kuo, C.-J., Chung, I., Hsu, C.-N., Lin, Y.-S., Klinger, R., Friedrich, C. M., Ganchev, K., Torii, M., et al. (2008). Overview of biocreative ii gene mention recognition. Genome biology, 9(2), 1–19.

Szklarczyk, D., Kirsch, R., Koutrouli, M., Nastou, K., Mehryary, F., Hachilif, R., Gable, A. L., Fang, T., Doncheva, N. T., Pyysalo, S., et al. (2023). The string database in 2023: protein–protein association networks and functional enrichment analyses for any sequenced genome of interest. Nucleic Acids Research, 51(D1), D638–D646.

The Uniprot Consortium (2021). Uniprot: the universal protein knowledgebase in 2021. Nucleic acids research, 49(D1), D480–D489.

Vaswani, A., Shazeer, N., Parmar, N., Uszkoreit, J., Jones, L., Gomez, A. N., Kaiser, L., and Polosukhin, I. (2017). Attention is all you need. In Proceedings of the 31st International Conference on Neural Information Processing Systems, NIPS’17, page 6000–6010, Red Hook, NY, USA. Curran Associates Inc.

Zhang, Y., Zhang, Y., Qi, P., Manning, C. D., and Langlotz, C. P. (2021). Biomedical and clinical English model packages for the Stanza Python NLP library. Journal of the American Medical Informatics Association, 28(9), 1892–1899.

